# SHORTENING TIME FOR ALCOHOL ACCESS DRIVES UP FRONT-LOADING BEHAVIOR, BRINGING CONSUMPTION IN MALE RATS TO THE LEVEL OF FEMALES

**DOI:** 10.1101/2021.06.01.446588

**Authors:** A. Flores-Bonilla, B. De Oliveira, A. Silva-Gotay, K. Lucier, H.N. Richardson

## Abstract

Alcohol can have more detrimental effects on mental health in women, even when intake is comparable or higher in men. This may relate to a differential pattern of drinking, e.g., how rapidly alcohol is consumed. We used operant procedures to gain insight into sex differences in the drinking dynamics of rats. Adult male and female Wistar rats underwent operant training to promote voluntary drinking of 10% (w/v) alcohol (8 rats/sex). We tested how drinking patterns changed after manipulating the effort required for alcohol (fixed ratio, FR), as well as the length of time in which animals had access to alcohol (self-administration session length). Rats were tested twice within the 12 hours of the dark cycle, at 2 hours (early sessions) and 10 hours into the dark cycle (late sessions). As expected, adult females consumed significantly more alcohol than males in the 30-minute sessions with the FR1 paradigm. Alcohol consumption within females was higher in the late sessions compared to early sessions, whereas this difference was not found within males. “Front-loading” of alcohol (heavier drinking in the first five minutes of the session) was the primary factor underlying higher consumption in females, and this sex difference was accentuated in the late sessions. Increasing the effort required from FR1 to FR3 reduced alcohol drinking in both sexes. Front-loading behavior remained in females in both early and late sessions, whereas males exhibited minimal front-loading behavior only in the early sessions. Compressing drinking access to 15-minutes drove up front-loading behavior, producing total alcohol intake levels that were comparable in both sexes. This strategy could be useful for exploring sex differences in the effect of voluntary alcohol drinking on the brain. Our findings also highlight the importance of the time of testing for detecting sex differences in drinking behavior.

**Highlights:**

- Voluntary alcohol drinking is higher in adult female rats compared to adult male rats. This sex difference is most pronounced in the later phase of the dark cycle, and when the operant effort is minimal (when 1 lever press gives 1 reward: fixed ratio 1, FR1).
- Higher alcohol intake in females is primarily due to “front-loading,” or the rapid consumption of alcohol within the first 5 minutes of access.
- Increasing the effort required to obtain alcohol from FR1 to FR3 dampens “front-loading” drinking behavior, resulting in similar levels of total intake in males and females.
- Compressing the time of access to 15 minutes drives up “front-loading” to such a degree that animals end up consuming more alcohol in total than they do in 30-minute sessions. In males, this increase in drinking is large enough that it eliminates the sex difference in total alcohol intake.

**Visual Abstract:** 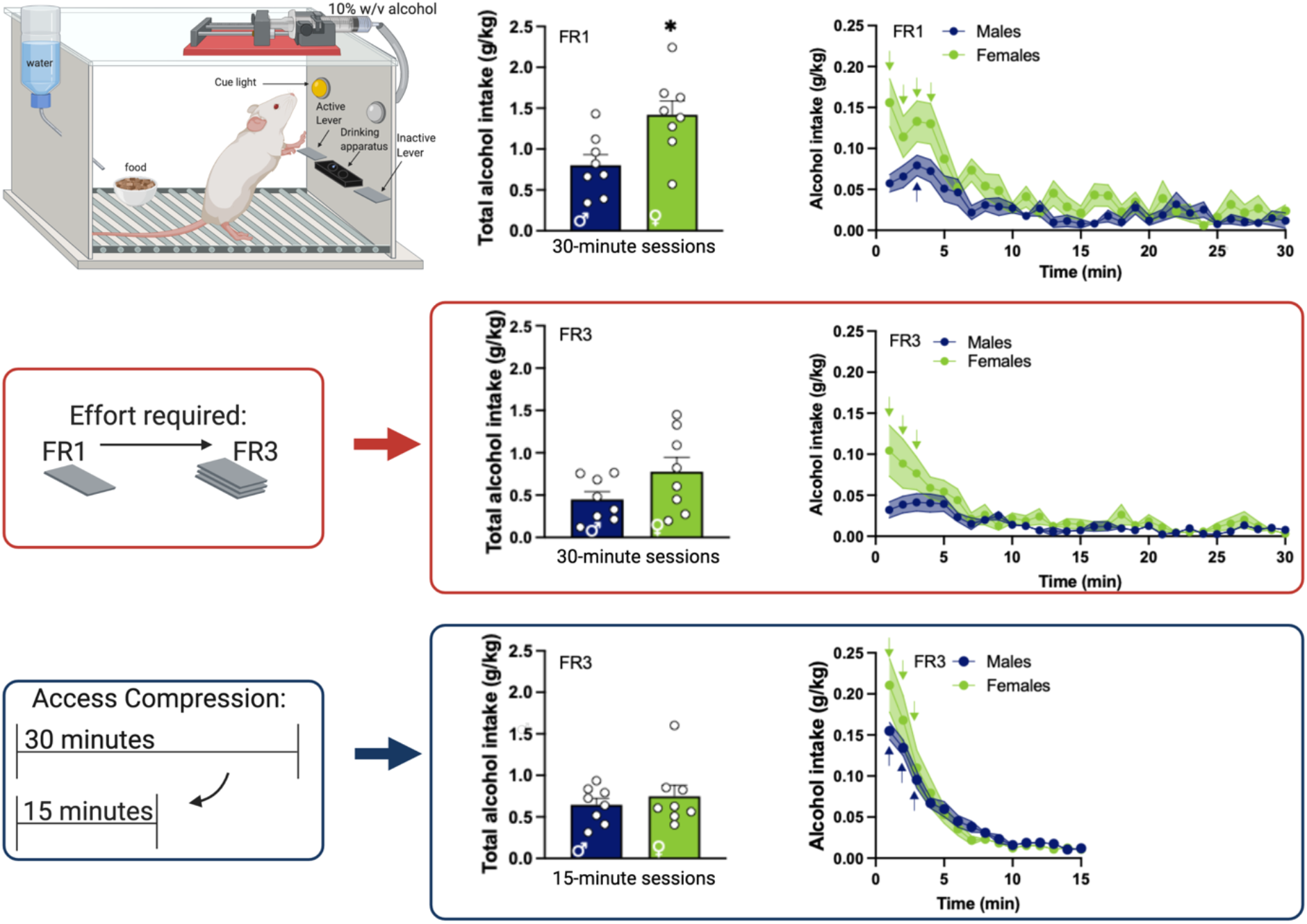

## 1. Introduction

Alcohol use disorder (AUD) is a chronic relapsing condition characterized by the misuse of alcohol (American Psychiatric Association, 2013; Grant et al., 2015; White et al., 2018). In the United States, 29% of adults have a lifetime prevalence for AUD, and men have a higher prevalence than women, yet this gap has been closing in recent years (Grant et al., 2017, 2015; White et al., 2017). Serious long-term effects of AUD are presented differently in men and women even when drinking levels are comparable, with some risks higher in women compared to men (Åberg et al., 2017; Hamajima et al., 2002; Hydes et al., 2019; Kerr-Corrêa et al., 2007; Loft et al., 1987; Mann et al., 2005; Schwarzinger et al., 2018; Smith-Warner et al., 1998; Urbano-Márquez et al., 1995; White et al., 2017); reviewed in (Erol and Karpyak, 2015; Peltier et al., 2019; Wilsnack et al., 2018). These negative consequences are exacerbated by excessive patterns of compulsive drinking; e.g., binge drinking (Fede et al., 2020; Kvamme et al., 2016; Mashhoon et al., 2014; National Institute on Alcohol Abuse and Alcoholism, 2004); reviewed in (Becker et al., 2012; George et al., 2014; Koob, 2013; Mason, 2017; Peltier et al., 2019). Binge drinking is considered the first of the three stages of addiction: Binge/Intoxication, Negative Affect/Withdrawal, and Preoccupation/Anticipation, with evidence of sex differences being reported in all three stages, reviewed in (Flores-Bonilla and Richardson, 2020). While the prevalence of binge drinking is higher in men (33%) than women (17%), women experience greater mental health instability (17.3% vs. 8.6%) and worse mental health consequences (11.2% vs. 6.3%) compared to men (Patrick et al., 2020; Substance Abuse and Mental Health Services Administration, 2014).

Operant self-administration in rodents is useful for modeling voluntary alcohol consumption in humans, while maintaining a level of experimental control needed for precise tracking and manipulation of alcohol drinking behavior (Brown et al., 1998; Elmer et al., 1987a, 1987b; George, 1987; Grant and Samson, 1986, 1985; June and Gilpin, 2010; Risinger et al., 1998); reviewed in (Green and Grahame, 2008; Jeanblanc et al., 2019). We used this approach to map out the dynamic microstructural patterns in drinking in male and female adult rats to explore various aspects of drinking behavior such as rapid drinking at the start of a session i.e., “front-loading” (Ardinger et al., 2020; Baird et al., 2005; D’Aquila et al., 2012; Davis, 1996; Lardeux et al., 2013; Linsenbardt and Boehm, 2015; Robinson and McCool, 2015). We also manipulated operant parameters of alcohol availability to further probe these sex differences in drinking patterns. We report herein that front-loading drives sex differences in alcohol intake, with greater front-loading behavior in females. Moreover, by manipulating operant parameters we were able to augment this rapid drinking pattern in males to reflect the levels observed in females.

## 2. Materials and Methods

### 2.1 Animals (subjects)

A total of 24 adult Wistar rats (12 per sex) were obtained from Charles River (Wilmington, MA, USA). Upon arrival, the animals were 85 days old and group-housed by sex (2 rats per cage) under a reverse light cycle of 12-hour light / 12-hour dark (lights OFF at 7 am / lights ON at 7 pm). Cages were standard plastic with wood chip bedding and animals had food and water access *ad libitum* with no deprivation. Animals acclimated for 15 days before operant training began. Rats were weighed once a day and weight data was used to calculate alcohol intake per weight (g/kg) of each rat. At the start of the experiment, males weighed 352.00 ± 5.42 g and females weighed 241.25 ± 4.81 g. The training sessions were conducted once a day and started after the first 2 hours of the dark cycle (at 9 am). The drinking sessions were conducted twice a day; the first session (“early”) started at 9 am (2 hours into the dark cycle) and the second session (“late”) started at 5 pm (10 hours into the dark cycle with an 8-hour break between sessions). All procedures were performed according to the National Institutes of Health Guide for the Care and Use of Laboratory Animals and approved by the Institutional Animal Care and Use Committee.

### 2.2 Voluntary Operant Self-Administration Experiments

#### 2.2.1 Five days of training sessions

Rats were randomly assigned to an alcohol group (16 rats; 8 per sex) or water control group (8 rats; 4 per sex) before the start of the training. On the first day of training, rats had two-bottle choice access to 10% (w/v) alcohol (ethanol) diluted in tap water or only tap water in their home-cage for 24 hours, to reduce novelty of the alcohol solution. A detailed timeline of experiments can be found in **Fig 1**. On the second day of training, rats were trained for 12 hours to orally self-administer water in operant boxes (Coulbourn Instruments, Allentown, PA). The operant boxes were individually housed inside ventilated cubicles to minimize noise and environmental disturbances. In each operant box, there were two retractable levers 4 cm above the grid floor and each lever was 4.5 cm on the side of a two-well acrylic drinking apparatus (custom made by Behavioral Pharma, San Diego, CA). One of the levers was active, which a response (lever press) consequently delivered 0.1 ml of fluid (water for all animals at this point of the experiment) from a 15 rpm Razel syringe pump (Stamford, CT) to the appropriate well over 0.5 seconds. The other lever was inactive, only recording the operant response without delivery of fluid. All lever presses and subsequent deliveries were recorded using custom Med-PC IV software. Both levers were available from the start of the session and did not retract until the end of the session. The active lever was programmed to a fixed ratio 1 (FR1) schedule and its location was counterbalanced between each rat (half of the animals had an active left lever and the other half had an active right lever). The stimulus light above the appropriate active lever for each rat was set to turn on every time the required number of presses for delivery was completed and served as a cue for fluid delivery in the well of the drinking apparatus. During the 0.5 seconds of pump activation, active lever presses were not recorded and did not result in the delivery of liquid. On the third day, rats remained in their home cage to rest. The next day, they were placed in the operant boxes for 2 hours of access to 10% (w/v) alcohol or tap water (0.1 ml delivered after each lever press). On the fifth day of training, rats were placed in the operant boxes and given 1-hour access to alcohol. After each operant session, the drinking apparatus was inspected for any leftover liquid to confirm animals had consumed all the alcohol (or water for control animals). Animals also had *ad libitum* access to food and water during all the operant sessions (a ceramic or stainless-steel dish containing rat chow and a water bottle were always available in each operant box). This ensured that lever presses for alcohol were not motivated by hunger or thirst. All data was collected as lever presses per minute of the session (1-minute bins) and as total lever presses per session.

**Figure 1.**
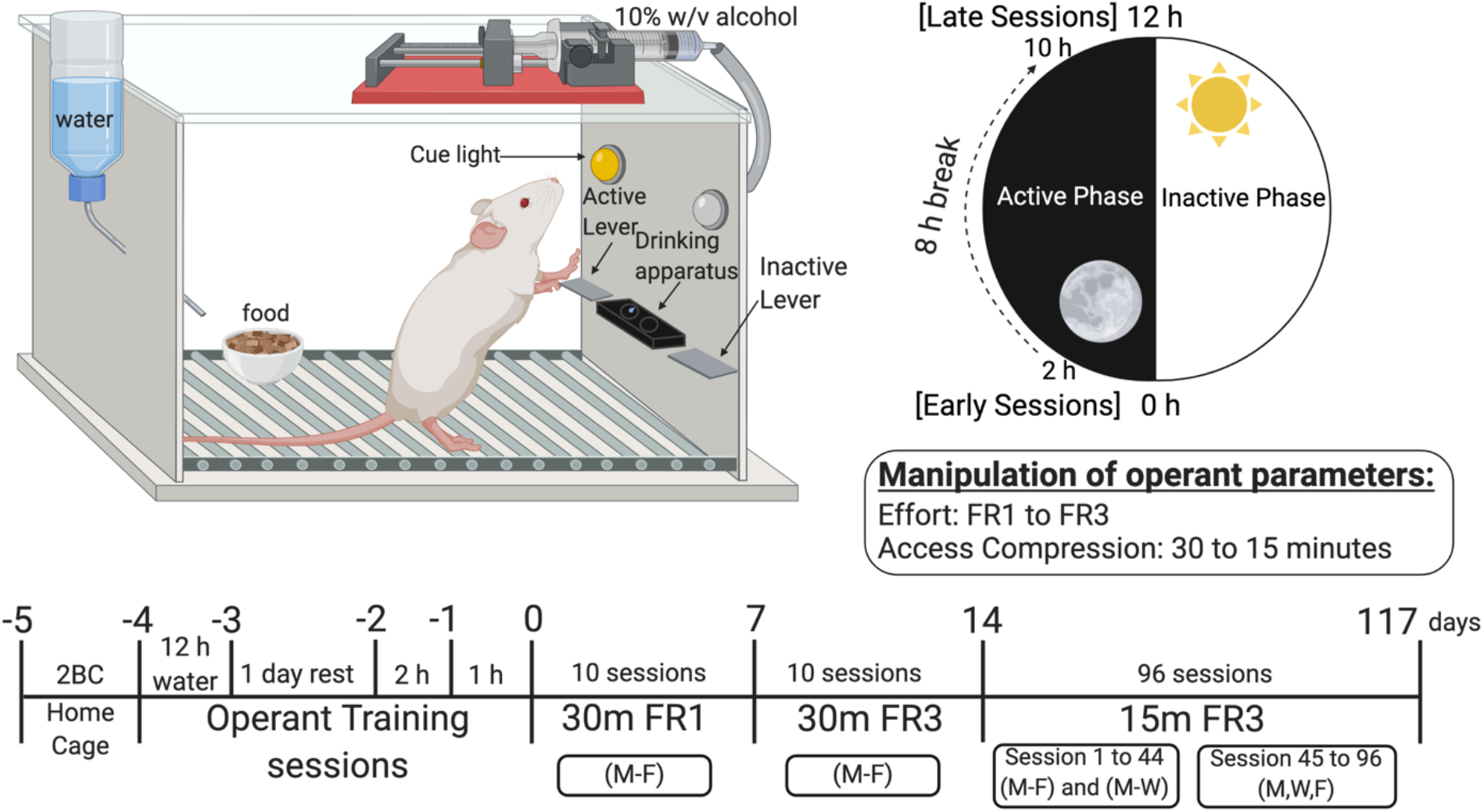
Operant Conditioning Boxes and Experimental Timeline. Male and female adult Wistar rats were trained to self-administer 10% w/v alcohol using operant conditioning. Training began with 1 day of 2BC in their home cages, followed by the first day of operant training for water (12 h session), a day of rest in the home cage, and two days of operant training for alcohol (2 h session on the first day and 1 h session on the second day). Operant testing was conducted twice each day, beginning 2 h (early session) and 10 h (late session) in the dark cycle for a total of 117 days. Operant parameters such as access time (30 minutes vs. 15 minutes), effort (fixed ratio, FR1 vs. FR3), and session time (early vs. late in the dark cycle) are summarized in the illustration and timeline. Operant boxes had two levers (one active and one inactive), as well as a food cup and water bottle that could be accessed *ad libitum.* Alcohol was delivered to a two-well acrylic drinking apparatus after every 1 press (FR1) or 3 presses (FR3). The cue light above the active lever is turned on for each alcohol delivery. *Note:* w/v, weight per volume; 2BC, two-bottle choice in the home cage; FR, fixed ratio; h, hour; m, minutes; M, Monday; W, Wednesday; F, Friday. Created with BioRender.com

#### 2.2.2 Manipulation of operant parameters

At the start of the drinking sessions the following day, rats were placed in the operant boxes twice a day with an 8-hour break between the sessions to quantify the alcohol intake difference between the early and late sessions. For the first week, rats had 30-minute access in an FR1 schedule for a total of 10 sessions or 5 days (Monday-Friday) and rested in their home cages on the weekend. The following Monday, operant parameters were changed to an FR3 schedule (1 delivery for every 3 presses) to increase effort for alcohol for the next 10 sessions (5 days, Monday-Friday). After rest on the weekend, the operant parameter of access time was compressed from 30 minutes to 15 minutes for 96 sessions or 48 days in total. For the first 20 sessions of the 15-minute FR3 schedule, rats were placed in the operant boxes five consecutive days of the week (Monday-Friday; continuous access) and had two rest days in their home cages on the weekend. Then, from session 21 to 44, rats were placed in the operant boxes for three consecutive days (Monday-Wednesday; continuous access) and had four consecutive rest days in their home cages (Thursday-Sunday). From sessions 45 to 96, rats were placed in the operant boxes for three alternating days (Monday, Wednesday, and Friday; intermittent access) and had four rest days (Tuesday, Thursday, Saturday, and Sunday). Analysis of these two different schedules showed no differences in total alcohol intake over the session (F _(1, 14)_ = 3.07, *p* > 0.05; data not shown) or in microstructural analysis per one-minute intake (F _(1, 14)_ = 0.01, *p* > 0.05; data not shown). Therefore, we pooled the data for analysis of the 15-minute FR3 sessions.

### 2.3 Statistical analysis

Statistical analyses and the generation of graphs were done using GraphPad Prism version 8.4.3 for Mac (GraphPad Software, San Diego, CA). Data are shown as the mean and standard error of the mean (± SEM) unless otherwise indicated. Total alcohol intake (g/kg), total water intake (ml/kg) and responding on the inactive lever (presses/min) data in each within-subject condition (normal effort vs. increased effort and normal access vs. compressed access) were analyzed using a two-way repeated-measures analysis of variance (RM-ANOVA) with session (early session vs. late session) as the within-subject factor and sex (male vs. female) as the between-subject factor. Bonferroni’s multiple comparisons test was conducted as *post hoc* analysis following significant main effects or interactions, as recommended (Armstrong, 2014; McHugh, 2011). For microstructural analysis, the 1-minute bins of drinking data were analyzed using a three-way, sex x session x time, with significant main effects and interactions further analyzed with two-way, sex x time RM-ANOVAs, with session and time (in 1-minute bins) as within-subject factors and sex as the between-subject factor. The significant main effects and interactions were followed up using Tukey’s multiple comparison *post hoc* tests to assess “front-loading” behavior in the first 5 minutes of access by comparing each 1-minute bin to the last 5 minutes of the session, as recommended when comparing simple effects (McHugh, 2011). Access compression was analyzed by three-way, sex x access compression x time and access compression x session x time, with significant main effects and interactions was further analyzed with two-way, access compression x time and sex x time RM-ANOVAs, with access compression and time as within-subject factors and sex as the between-subject factor. Bonferroni’s multiple comparisons test was conducted as *post hoc* analysis following significant main effects or interactions, by comparing the total intake in each 1-minute bin between the normal access and the compressed access sessions. To determine whether total alcohol consumed in the first 5-minutes differed by sex, we used unpaired two-tailed t-tests. The accepted level of significance for all tests was p ≤ 0.05 and was indicated by different symbols (summarized in **Table 1**).

**Table 1.** Overview of statistical indicators used in the figures.

| Symbol | Significant effect | Statistical analysis | Used in |
| --- | --- | --- | --- |
| * | Sex differences | Bonferroni's multiple comparisons *post hoc* test | Fig. 2A, Inset of Fig. 2D |
| # | Effect of session | Bonferroni's multiple comparisons *post hoc* test | Fig. 2A, 3A |
| ↓ | Front-loading | Tukey's multiple comparisons *post hoc* test | Fig. 2C, 2D, 3C, 3D, 4C, 4D, 4E, 4F |
| + | Access compression | Bonferroni's multiple comparisons *post hoc* test | Fig. 4A, 4C, 4D, 4E, 4F |

## 3. Results

### 3.1 Front-loading drives sex differences in voluntary alcohol intake and is greatest when testing occurs later in the dark cycle

Sex differences in alcohol consumption emerge by adulthood in rodents, with females drinking more than males (Juárez and De Tomasi, 1999; Randall et al., 2017); for review see (Flores-Bonilla and Richardson, 2020). As expected, the average of total alcohol intake (g/kg) within 30-minute sessions was higher in adult female rats compared to males (**Fig. 2A**; main effect of sex, F _(1, 14)_ = 6.67, *p* < 0.05). Alcohol intake was higher in the late sessions compared to the early sessions (main effect of session, F _(1,14)_ = 34.97, *p* < 0.0001). There was also a session x sex interaction (F _(1,14)_ = 6.34, *p* < 0.05) and *post hoc* analysis showed significantly higher alcohol intake in females compared to males only in the late sessions (**Fig. 2A**; Bonferroni’s multiple comparisons test, adjusted *p* < 0.01), indicating that sex differences in alcohol drinking were greatest when testing was conducted later in the dark cycle and not earlier in the dark cycle (adjusted *p* > 0.05). Rats consumed significantly more alcohol in the late sessions compared to the early sessions within females (**Fig. 2A**; adjusted *p* < 0.0001), whereas within males, this was only a trend (adjusted *p* = 0.06). Higher alcohol intake in the later sessions was not due to differential levels of non-specific activity, as inactive lever presses were comparable in the early and late sessions (F _(1, 14)_ = 0.06, *p* > 0.05; **Table 2**). In control animals, self-administration of water did not differ between males and females (**Fig. 2B**; F _(1, 6)_ = 0.61, *p* > 0.05), or between the early and late sessions (F _(1, 6)_ = 1.35, *p* > 0.05).

**Figure 2.**
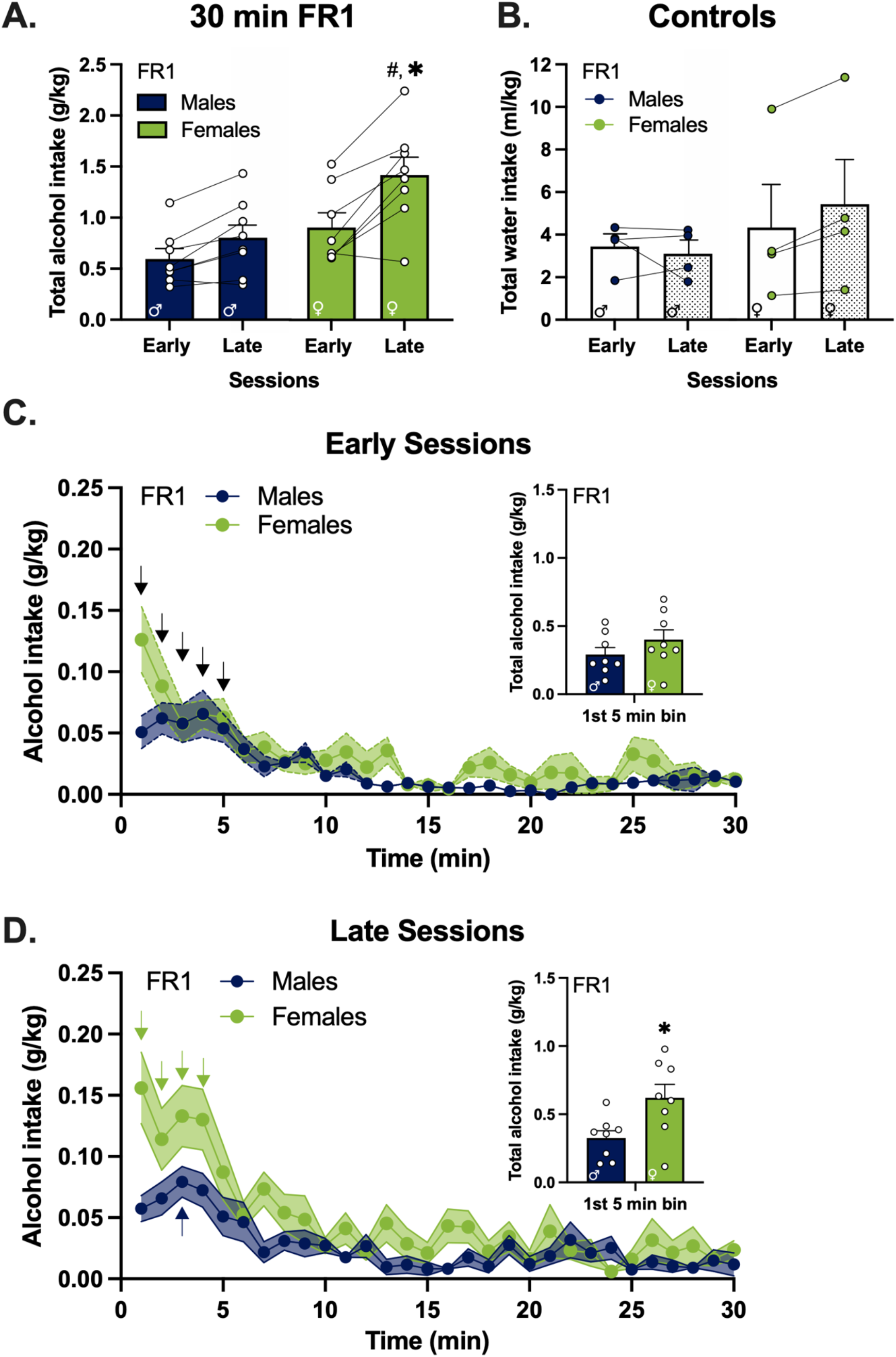
Front-loading drinking behavior drives sex differences in alcohol intake on the sessions late into the dark cycle. **A)** Average of total alcohol intake of early and late sessions (n = 8 / sex); **B)** Average of total water intake in the control group (n = 4 / sex); **C)** Early sessions and **D)** late sessions of total alcohol intake per minute in the session, inset: total alcohol intake in the first 5-minute bin. *Note:* FR, fixed ratio. **A)** #, higher intake in late sessions compared to early sessions in females, adjusted *p* <0.05, and **\***, higher intake in females compared to males in the late sessions, adjusted *p*<0.05; both indicated by Bonferroni’s multiple comparisons test *post hoc* analysis following a session x sex interaction. **C)** black arrows, higher intake compared to each minute of the last 5 minutes of the session collapsed across sex, adjusted *ps* < 0.05; indicated by Tukey’s multiple comparisons test *post hoc* analysis following a main effect of time. **D)** arrows, higher intake compared to each minute of the last 5 minutes of the session in females (green arrows) and males (blue arrow), adjusted *ps* < 0.05; indicated by Tukey’s multiple comparisons test *post hoc* analysis following a time x sex interaction. In inset graph of **D) \***, unpaired t-test, *p*<0.05. Error bars and shading are shown in SEM.

**Table 2.** Effect of sex and session on inactive lever presses.

| Operant parameters | Inactive lever (presses/minute)<br>Mean (± SEM) | | | RM ANOVA: sex x session | | | |
| --- | --- | --- | --- | --- | --- | --- | --- |
|  | Session | Males | Females | Effect | *p*-value | F | Df |
| 30-minute<br>FR1<br>sessions | **Early** | 0.2044<br>(0.03) | 0.1506<br>(0.03) | **Sex** | 0.9536 | 0.003 | 1, 14 |
|  | **Late** | 0.1500<br>(0.03) | 0.1844<br>(0.03) | **Session** | 0.8154 | 0.06 |  |
|  |  |  |  | **Sex x Session** | 0.1623 | 2.18 |  |
| 30-minute<br>FR3<br>sessions | **Early** | 0.1039<br>(0.02) | 0.0806<br>(0.01) | **Sex** | 0.2615 | 1.37 | 1, 14 |
|  | **Late** | 0.0872<br>(0.02) | 0.1511<br>(0.03) | **Session** | 0.0620 | 4.12 |  |
|  |  |  |  | **Sex x Session** | 0.0780 | 3.62 |  |
| 15-minute<br>FR3<br>sessions | **Early** | 0.0703<br>(0.02) | 0.0571<br>(0.01) | **Sex** | 0.7718 | 0.09 | 1, 14 |
|  | **Late** | 0.0839<br>(0.03) | 0.0764<br>(0.01) | **Session** | 0.0114<sup>#</sup> | 8.48 |  |
|  |  |  |  | **Sex x Session** | 0.4882 | 0.51 |  |

RM-ANOVA = repeated measures analysis of variance; <sup>#</sup>p<0.05

We followed with a microstructural analysis of alcohol drinking, measuring the total alcohol intake per minute across the operant session. In the early sessions, time had a significant effect on alcohol intake (**Fig. 2C**; main effect of time, F _(29, 406)_ = 10.40, *p* < 0.0001) but there were no differences between males and females (F _(1,14)_ = 3.81, *p* = 0.07). *Post hoc* analyses were therefore collapsed across sex. Both males and females showed significantly higher alcohol intake in the first 5 minutes of the session compared to the alcohol intake in each minute of the last 5 minutes of the session (Tukey’s multiple comparisons test, adjusted *p*s < 0.05) which is characteristic of “front-loading” (black arrows, **Fig. 2C**). In contrast, animals showed a significant time x sex interaction (**Fig. 2D**; F _(29, 406)_ = 2.07, *p* < 0.001) in the late sessions, with females showing higher alcohol intake in the first 4 minutes of the session compared to the last 5 minutes of the session (Tukey’s multiple comparisons test, adjusted *p*s < 0.05; green arrows in **Fig. 2D**). In males, this front-loading drinking behavior was transiently observed (only minute 3 had significantly higher intake compared to the last 5 minutes of the session, Tukey’s multiple comparisons test, adjusted *p*s < 0.05; blue arrow in **Fig. 2D**). The total amount of alcohol ingested during front-loading was higher in females compared to males in the late sessions (**Fig. 2D inset**; unpaired t-test, p < 0.05), but not in the early sessions (**Fig. 2C inset**; unpaired t-test, *p* > 0.05).

### 3.2 Increasing effort suppresses sex differences in total alcohol intake

To test if increasing the effort to obtain alcohol reduces drinking to the same degree in both males and females, we manipulated the fixed ratio schedule of deliveries from FR1 to FR3, i.e., shifting from requiring 1 to requiring 3 presses for alcohol delivery. This increase in effort suppressed the total amount of alcohol self-administered during the 30-minute session in both male and female rats (main effect of effort, F _(1, 14)_ = 27.87, *p* = 0.0001 in early sessions and F _(1, 14)_ = 31.10, *p* < 0.0001 in late sessions), resulting in a loss of sex difference in total alcohol intake (**Fig. 3A**; F _(1, 14)_ = 2.22, *p* > 0.05; for comparisons to FR1, see **Fig 2**). There was a main effect of session (**Fig. 3A**; F _(1, 14)_ = 14.96, *p* < 0.01) and a session x sex interaction (**Fig. 3A**; F _(1,14)_ = 4.43, *p* = 0.05). *Post hoc* analysis showed higher alcohol consumption in the late sessions compared to the early sessions within females (Bonferroni’s multiple comparisons test, adjusted *p* < 0.01), but not within males. In the FR3 sessions, control animals continue to show a lack of sex differences in self-administration of water (**Fig 3B**; F _(1,6)_ = 0.37, *p* > 0.05). There was also no effect of time of session on self-administration of water (**Fig 3B**; F _(1,6)_ = 1.31, *p* > 0.05). There was a trend for higher responding on the inactive lever later in the dark cycle, but this was not significant (F _(1, 14)_ = 4.12, *p* = 0.062), along with a non-significant trend for a session x sex interaction (F _(1, 14)_ = 3.62, *p* = 0.078; shown in **Table 2**).

**Figure 3.**
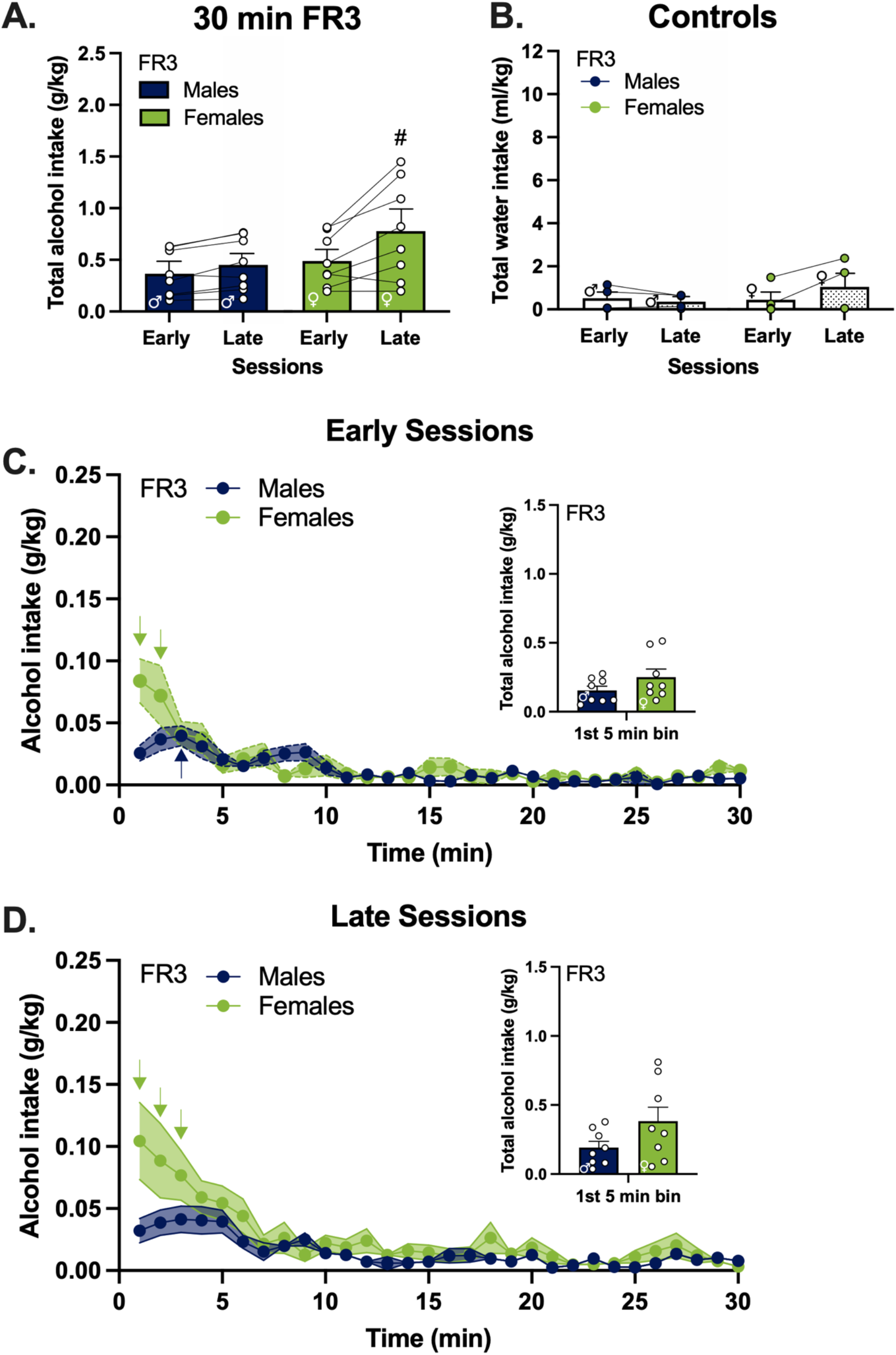
Females maintain front-loading behavior after increasing effort in the late sessions despite an overall decrease in total alcohol drinking. **A)** Average of total alcohol intake of early and late sessions (n = 8 / sex); **B)** Average of total water intake in the control group (n = 4 / sex); **C)** Early sessions and **D)** late sessions of total alcohol intake per minute in the session, inset: total alcohol intake in the first 5-minute bin. *Note:* FR, fixed ratio. **A)** #, higher intake in the late sessions compared to the early sessions in females, adjusted *p* < 0.01; indicated by Bonferroni’s multiple comparisons test *post hoc* analysis following a main effect of session. **C), D)** arrows, higher intake compared to each minute of the last 5 minutes of the session in females (green arrows) and males (blue arrow), adjusted *ps* < 0.05; indicated by Tukey’s multiple comparisons test *post hoc* analysis following a time x sex interaction. Error bars and shading are shown in SEM.

### 3.3 Females exhibit higher levels of front-loading drinking behavior compared to males

We next tested whether sex differences in the microstructural analysis of drinking persisted when animals had to work harder for alcohol. In the early sessions, there was a main effect of time (**Fig. 3C**; F _(29, 406)_ = 10.75, *p* < 0.0001) and time x sex interaction (F _(29, 406)_ = 2.44, *p* < 0.0001), and *post hoc* analysis showed higher alcohol intake in the first 2 minutes of the session compared to the last 5 minutes of the session in females (Tukey’s multiple comparisons test, adjusted *p*s < 0.05; green arrows in **Fig. 3C**) and only in minute 3 of the session in males compared to the last 5 minutes of the session (Tukey’s multiple comparisons test, adjusted *p*s < 0.05; blue arrow in **Fig. 3C**). In the late sessions there was a main effect of time (**Fig. 3D**; F _(29, 406)_ = 10.59, *p* < 0.0001) and time x sex interaction (F _(29, 406)_ = 2.18, *p* < 0.001), and in contrast to the early sessions, *post hoc* analysis showed higher alcohol intake in the first 3 minutes of the session compared to the last 5 minutes of the session in females (Tukey’s multiple comparisons test, all adjusted *p*s < 0.05; green arrows in **Fig. 3D**), while males showed no significant differences (Tukey’s multiple comparisons test, adjusted *p*s > 0.05). This suggests that under FR3 conditions, females still exhibit more front-loading behavior compared to males, especially later in the dark cycle. However, despite higher levels of front-loading in females, the total amount of alcohol consumed in the first 5 minutes of the session was no longer significantly different between males and females in the early or late sessions (**Fig. 3C inset**, early sessions, and **3D inset**, late sessions; unpaired t-test, *p* > 0.05).

### 3.4 Compressing access time augments front-loading drinking behavior resulting in more total consumption of alcohol

To determine whether reducing the time of operant sessions affects front-loading behavior in both males and females, we reduced alcohol access in the sessions from 30 minutes to 15 minutes (access compression). Limiting alcohol availability elicited higher front-loading drinking behavior in both males and females in the early sessions, and this rapid rate of drinking resulted in more total alcohol intake overall (**Fig. 4A**; main effect of access compression, with the shorter sessions eliciting higher total intake compared to the longer sessions, F _(1, 14)_ = 28.65, *p* = 0.0001). In the late sessions, access compression had no significant effect (**Fig. 4B**; F_(1, 14)_ = 1.44, *p* > 0.05) as animals of both sexes consumed similar amounts of alcohol despite having only half the time to drink. These data altogether indicate that limiting access causes animals to voluntarily drink more quickly when giving the opportunity, more than doubling drinking levels in the first few minutes.

**Figure 4.**
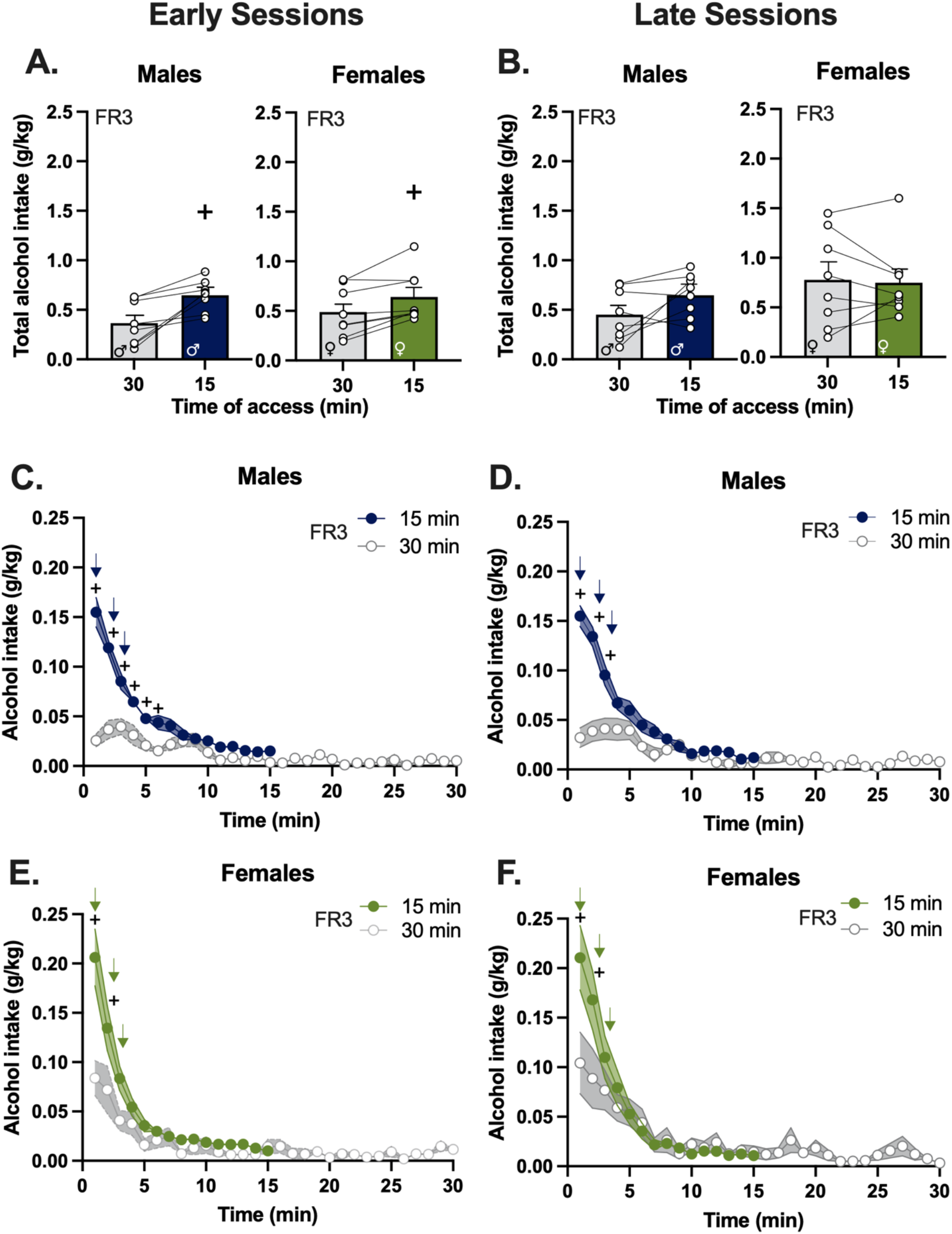
Access time compression increased front-loading of alcohol intake of males to the level of intake in females. **A)** Average of total alcohol intake of early and **B)** late sessions (n = 8 / sex); **C)** Early sessions and **D)** late sessions of total alcohol intake per minute in the session of males; **E)** Early sessions and **F)** late sessions of total alcohol intake per minute in the session of females. *Note:* FR, fixed ratio. **A)** +, higher intake in the compressed access (15-minute) sessions compared to the normal (30-minute) sessions in both males and females, adjusted *p* < 0.05; indicated by a main effect of access compression. **C-F)** +, higher intake in the compressed access (15-minute) sessions compared to the normal (30-minute) sessions in males (**C** and **D**) and females (**E** and **F**), adjusted *p*s < 0.05; indicated by Bonferroni’s multiple comparisons test *post hoc* analysis following time x access compression interaction. **C-F)** arrows, higher intake compared to each minute of the last 5 minutes of the session in females (green arrows) and males (blue arrows), adjusted *p*s < 0.05; indicated by Tukey’s multiple comparisons test *post hoc* analysis following time x sex interaction. Error bars and shading are shown in SEM.

To gain insight into the specific patterns of drinking that lead to higher total intake in the compressed sessions, we compared the alcohol intake in 1-minute bins of the compressed sessions with the normal (30-minute) sessions. We found that compression of access time increased front-loading of males in the early sessions (**Fig. 4C**; time x access compression interaction, F _(14, 196)_ = 24.94, *p* < 0.0001) and late sessions (**Fig. 4D**; time x access compression interaction, F _(14, 196)_ = 24.06, *p* < 0.0001). In females, compression of access time also increased front-loading in the early sessions (**Fig. 4E**; time x access compression interaction, F _(14, 196)_ = 5.90, *p* < 0.0001), and in the late sessions (**Fig. 4F**; time x access compression interaction, F _(14, 196)_ = 3.99, *p* < 0.0001). *Post hoc* analyses of the early sessions showed significantly higher alcohol intake after access compression in the first 6 minutes of the early sessions and the first 3 minutes of the late sessions in males of the compressed sessions compared to the normal sessions (+s in **Fig. 4C** and **Fig. 4D**, Bonferroni’s multiple comparisons test, adjusted *p*s < 0.05). In females, *post hoc* analyses showed significantly higher alcohol intake in the first 2 minutes in both early and late sessions of the compressed sessions compared to the normal sessions (+s in **Fig. 4E** and **Fig. 4F**, Bonferroni’s multiple comparisons test, adjusted *p*s < 0.05).

Between males and females, alcohol intake in 15 minutes of access time showed a significant time x sex interaction (F _(14, 196)_ = 2.31, *p* < 0.01) in the early sessions, as well as in the late sessions (F _(14, 196)_ = 2.20, *p* < 0.01). *Post hoc* analyses indicated that sex differences in drinking were only found in the first minute of the access in both early and late sessions (Bonferroni’s multiple comparisons test, adjusted *p*s < 0.05; these subtle sex differences can be viewed visually comparing the first minute in **Fig 4C** versus **Fig 4E** and **Fig 4D** versus **Fig 4F**). Thus, access compression drives up front-loading behavior to such an extent in males that alcohol drinking is almost identical to what is observed in females. Furthermore, both males and females consume significantly more alcohol in the first 3 minutes of the session compared to the last 5 minutes of the session (Tukey’s multiple comparisons test, adjusted *p*s < 0.05; arrows in **Fig. 4 C, D, E, F**) in both early and late sessions.

Control animals showed no differences in water intake between males and females (F _(1, 6)_ = 1.77, *p* > 0.05) nor differences between time of the session (F _(1, 6)_ = 0.05, *p* > 0.05, data not shown). The non-specific activity was found higher in the late sessions compared to the early sessions by an increase of inactive lever presses per minute (main effect of session, F _(1, 14)_ = 8.48, *p* < 0.01; shown in **Table 2**) with *post hoc* analysis showing significant differences in males (Bonferroni’s multiple comparisons test, adjusted *p*s < 0.05), but not females. This significant effect was not found in the active lever presses, as measured by the total alcohol intake (F _(1, 14)_ = 2.97, *p* > 0.05; data not shown).

## 4. Discussion

### 4.1 Main findings

The factors motivating drinking and the health consequences of heavy alcohol use can differ with sex (Flores-Bonilla and Richardson, 2020). Adolescent girls report drinking alcohol as a coping mechanism to alleviate depression and psychological distress because of its acute anxiety-reducing properties, while drinking in adolescent boys tends to relate more to risk-taking and impulsive behaviors (Bekman et al., 2013; Kuntsche and Müller, 2012). Ultimately in adulthood, women experience greater detrimental emotional and physical consequences including emotional instability, depression, and anxiety than men who engage in binge drinking (Orio et al., 2018; Patrick et al., 2020). Thus, there is a need to study not only the motivators that can lead to moderate or heavy alcohol use, but also how differential patterns of drinking behavior may influence the severity of negative consequences of alcohol on the brain. These questions can be answered using rodent models; however, it can be challenging to test the effect of alcohol drinking on the nervous system and behavior in rodents because adult female rodents tend to drink significantly more g/kg alcohol than their male counterparts (Juárez and De Tomasi, 1999; Lancaster and Spiegel, 1992). While this is well known in the field, the specific differences in patterns of drinking between males and females and their effects on the brain are not well understood. Herein, we used operant procedures to characterize the temporal dynamics of alcohol drinking in adult male and female rats to identify the specific pattern of drinking that drives the sex differences in intake and tested how changes in alcohol access can influence these sex differences. Collectively, we found that higher intake in females occurs during the initiation of access (i.e., “front-loading”). Moreover, by manipulating operant conditions such as access time, we were able to drive up front-loading behavior in males to levels that are comparable to females. This study overall provides insight into sex differences in drinking behavior, and also provides a novel approach to control voluntary intake in order to study its effects on the brain.

### 4.2 Front-loading drives sex differences in alcohol intake

Consistent with the characterization of sex differences in voluntary alcohol intake in rodents, female rats in our study showed higher total alcohol intake than males in the 30-minute sessions with minimal effort (Cofresí et al., 2019; Crabbe et al., 2009; Juárez and De Tomasi, 1999; Lancaster and Spiegel, 1992; Mankes et al., 1991; Randall et al., 2017); reviewed in (Flores-Bonilla and Richardson, 2020; Lancaster and Spiegel, 1992). Alcohol intake peaked in the first 5 minutes of the session, which provides evidence of front-loading behavior similar to other studies that identified higher intake of rewarding liquids (including alcohol in rats exposed to alcohol vapors, intermittent access to high-fat sweetened liquid in rats, and NaCl solution in sodium-depleted mice) at the start of access (D’Aquila et al., 2012; Darevsky et al., 2018; Davis, 1996; Lardeux et al., 2013; Linsenbardt and Boehm, 2015; Robinson and McCool, 2015; Siciliano et al., 2019). Additionally, front-loading can be suppressed with conditioned taste aversion with a bitter solution (LiCl) in the first 9 minutes of access (Baird et al., 2005). Reward expectation can induce front-loading to non-rewarding solutions in alcohol-exposed mice that receive water when alcohol is expected (Ardinger et al., 2020). Increasing the effort to obtain alcohol with an FR3 schedule suppressed the total alcohol intake in both males and females; however subtle sex differences in front-loading remained, as females showed front-loading while it was notably reduced in males. Compressing the access to alcohol eliminated the sex differences by increasing the front-loading of males to the level of females. As such, this data provides both insights into how microstructural patterns of drinking differ with sex and highlights an effective experimental strategy to effectively eliminate sex differences in voluntary drinking in adult rats.

### 4.3 Manipulation of operant parameters as a strategy to study sex differences

One challenge that researchers face when testing the effects of voluntary alcohol drinking on the brain is that differential levels of alcohol intake between males and females can complicate the experimental design. To circumvent this sex difference, involuntary/forced models of alcohol administration can be used which allows for delivery of equal amounts of low, moderate, and high levels of alcohol in males and females, albeit with different degrees of aversive consequences; for a review of current preclinical models of alcohol drinking see (Jeanblanc et al., 2019). In voluntary drinking models, operant parameters can be manipulated to bring one group’s intake down to the level of another group. For example, operant lever access can be controlled to limit the maximum number of rewards to match glucose intake in control groups consuming sweetened water to the glucose intake in groups consuming sweetened alcohol (Gilpin et al., 2012; Karanikas et al., 2013; Vargas et al., 2014). Conceptionally this same strategy could be used to reduce alcohol intake in adult females to the level of adult males, but this limits experimental design to testing only lower levels of voluntary alcohol. Herein, we were able to adjust the operant parameters to lead to higher alcohol intake in males, eliciting female-like drinking patterns; thus, increasing opportunities for testing the effects of voluntary alcohol on the brain and behavior. Furthermore, this rodent model of voluntary alcohol drinking can be used to study the neuronal activity underlying front-loading behavior, which ultimately may help develop targeted treatment strategies for alcohol abuse and AUD.

### 4.4 Effect of time of day on alcohol intake

Alcohol drinking tests in rodents are often conducted within the first few hours of the dark cycle to take advantage of increased nocturnal activity (Lei et al., 2016; Morales et al., 2015; Rhodes et al., 2005; Szumlinski et al., 2019; Thiele et al., 2014); reviewed in (Thiele and Navarro, 2014). In contrast, we show that alcohol consumption within females was highest when testing was done 10 hours, compared to 2 hours, into the dark cycle. This was not likely an effect of non-specific activity, because inactive lever presses did not differ between early and late sessions in females. This suggests that there is another factor that is driving up drinking in females when testing is done later in the dark cycle. It should be noted that, due to the within-subjects design of the study, all rats were tested twice a day; in which early sessions may act as a primer for increased alcohol drinking in the late sessions. To explore this further, we used a data set from a day in which animals only had a late session due to a power outage earlier in the day, which had to be excluded from the overall statistical analysis. We compared the drinking behavior of that day to a single late session from two days before (in which animals had an early session as well) and found no differences in alcohol intake. This suggests that those early sessions are not necessary for driving higher levels of intake in the late sessions. It is more likely that factors that change with circadian rhythms drive higher drinking in the later sessions. Indeed, baseline plasma corticosterone levels in rodents have a temporally regulated rhythm, with higher levels peaking around two hours before the start of the dark cycle and gradually decreasing until reaching low levels around 10 hours into the dark cycle in both males and females (Atkinson and Waddell, 1997; Yoshida et al., 2005). Low corticosterone levels in rodents and low cortisol levels in humans have been associated with higher drinking (Anthenelli et al., 2018; Orio et al., 2018; Richardson et al., 2008), reviewed in (Lu and Richardson, 2014). Notably, the circadian variation of plasma corticosterone levels is not found in *Crh*-deficient mice, suggesting that this is mediated by corticotropin-releasing factor signaling (Yoshida et al., 2005). Future studies could help determine whether circadian regulation play a role in the observed sex differences in late session alcohol intake.

### 4.5 Blood alcohol levels

Front-loading behavior induces most of the alcohol drinking in the first 5 minutes of the session, regardless of the total length of the session (30 vs. 15 minutes). Based on our current findings on front-loading, this suggests that collection of blood samples should take place after a few minutes of initial alcohol access for accuracy in binge drinking rodent models. Blood alcohol levels were not measured in this study; however, based on previous studies in male rats, the total consumption of 1.25 g/kg of alcohol or more reaches the blood alcohol levels of 0.08 g/dL or higher (Gilpin et al., 2012). However, this measurement was not evaluated in females. Other studies that use the “Drinking in the Dark” paradigm show that adult female mice consume more alcohol than males; however, these mice failed to consistently replicate the blood alcohol concentration levels that characterize binge drinking; i.e., 0.08 g/dL or higher (Szumlinski et al., 2019). Blood alcohol concentration levels are usually taken after testing typically around 30 minutes to 1 hour after the first initial intake (Carnicella et al., 2009); reviewed in (Jeanblanc et al., 2019). Female rodents metabolize alcohol at a greater rate than males while the opposite is true in humans (Teschke and Wiese, 1982); reviewed in (Jeanblanc et al., 2019; Müller, 2006). An important goal for future studies is to assess more accurate blood alcohol levels early in the operant session in both male and female rats while also taking into account the detrimental effects of alcohol metabolites in the brain (Baliño et al., 2019); reviewed in (Erol and Karpyak, 2015; Israel et al., 2015).

## 5. Conclusion

Sex differences are increasingly prevalent for studying alcohol binge drinking and AUD. Understanding the underlying patterns of drinking behaviors that drive these differences may help in determining factors for increased vulnerability for developing AUD. Overall, manipulation of operant parameters of alcohol availability altered behavioral patterns of drinking in both males and females. Manipulation of these parameters can be used to effectively study sex differences in alcohol-driven effects on the brain. Front-loading behavior is highest 10 hours into the dark cycle in females which drives the sex differences in the total alcohol intake. However, under certain conditions, adult male rats drink as much alcohol as females. Ultimately, the time of access to alcohol, the effort required to obtain it, and the time of day of testing all contribute to changes in front-loading behavior and increasing or decreasing the magnitude of sex differences in voluntary alcohol intake in adult rats.

## Acknowledgments

We thank the contribution of Dr. Said Akli, Lynn Bengtson, Sirisha Nouduri, Rithika Senthilkumar, Jillian Davis, and other members of the Richardson laboratory for critical feedback on data analysis, writing, and graphs. We also like to thank undergraduate students William Collins, Madeleine Pereira, Connor Murphy, and Eliza Kola for assistance with the behavioral experiments. Funding comes from the National Institutes of Health Postbaccalaureate Research Education Program grant NIH R25GM08624 that funded the participation of Annabelle Flores-Bonilla in the program, the National Science Foundation Graduate Research Fellowship Program NSF Award #1938059 to Annabelle Flores-Bonilla, NIH F99/K00 grant 1F99NS115272-01 to Andrea Silva-Gotay, and the National Institute on Alcohol Abuse and Alcoholism grant R01AA024774 to Dr. Heather N. Richardson.

## Author’s Contributions

A.F.B, K.L., and H.N.R. designed research; A.F.B., B.D., and K.L. performed research; A.F.B. and A.S.G. analyzed data; A.F.B, B.D., A.S.G, and H.N.R. wrote the paper.

